# High-Performance Wide-Band Open-Source System for Acoustic Stimulation

**DOI:** 10.1101/2024.02.22.581424

**Authors:** Artur Silva, Filipe Carvalho, Bruno F. Cruz

**Author notes:** The two authors contributed equally to this work (Artur Silva, Filipe Carvalho). Allen Institute for Neural Dynamics, Seattle, United States (Bruno F. Cruz).

## Abstract

The design and characterization of a low-cost, open-source auditory delivery system to deliver high performance auditory stimuli is presented. The system includes a high-fidelity sound card and audio amplifier devices with low-latency and wide bandwidth targeted for behavioral neuroscience research. The characterization of the individual devices and the entire system is performed, providing a thorough audio characterization data for varying frequencies and sound levels. The system implements the open-source Harp protocol, enabling the hardware timestamping of devices and seamless synchronization with other Harp devices.

**Specifications table:** 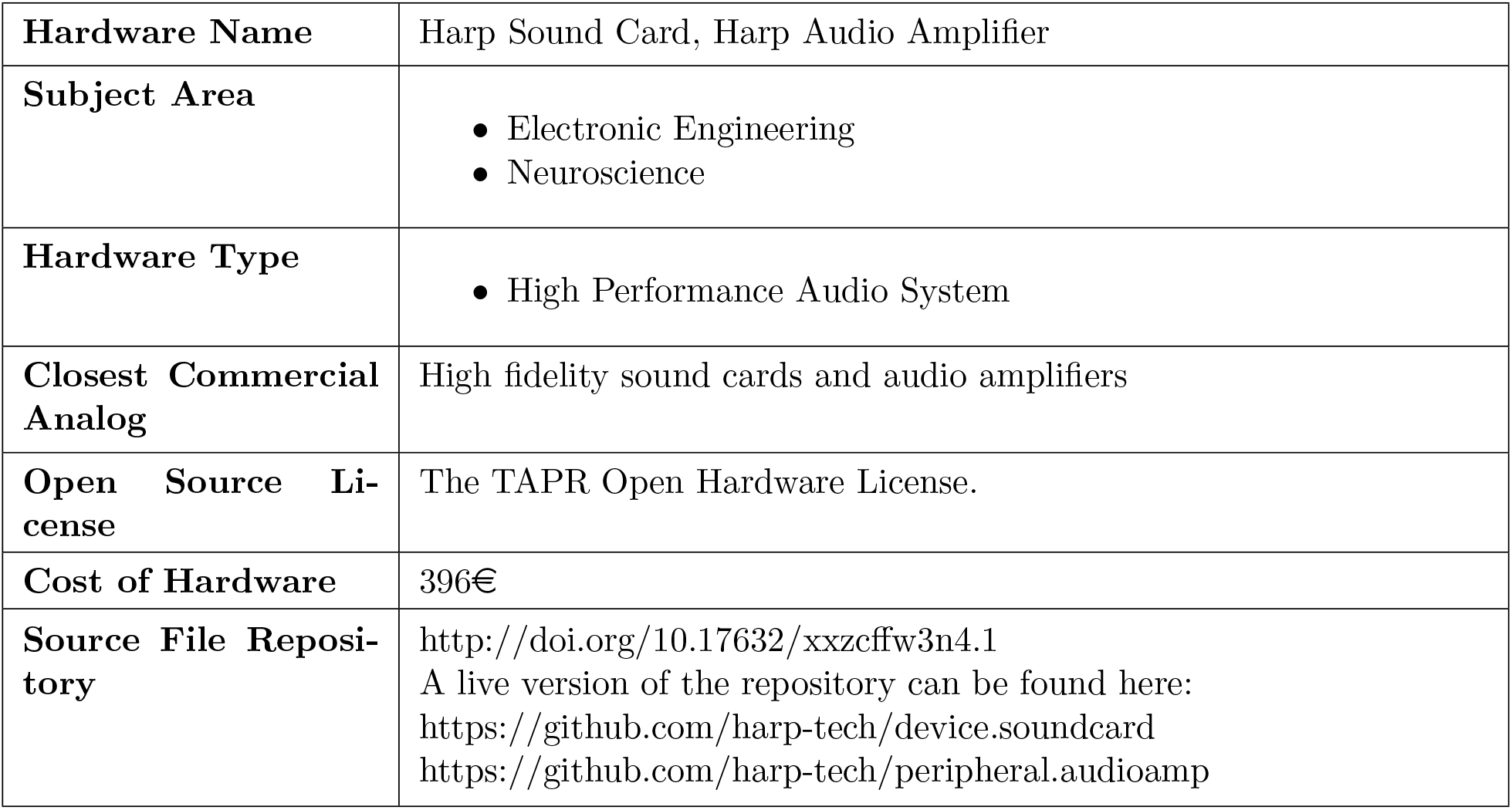

## 1. Hardware in context

Experimental behavior neuroscience relies on the ability to observe and measure the environment the animal lives in, as well as manipulate it. For most animals, and especially mammals, hearing is a critical sense. Among others, sound pressure levels can be used to provide critical information about where to forage or hunt, detect and avoid threats, navigate, and even communication with con specifics. While much can be learned from recording auditory features from the environment, it is often also necessary to “close the loop” by generating experimentally controlled stimuli.

Sound can be defined as a longitudinal pressure wave that travels in both space and time through a physical medium. An amplitude and single frequency characterize pure tone waveforms. As one might expect, the domain of these features in which animals can sense and produce sound partially defines their umwelt [12] and are distinct across species. For instance, while Human hearing is sensitive to frequencies of 30Hz-20kHz, mice expand the higher range to more than 85kHz [6].

Unfortunately, devices capable of generating high quality and experimentally controlled audio stimuli across a large section of this rich feature space are not widely available. Consumer-grade high-fidelity audio cards and amplifiers, while generally accessible and easy to use, are designed to operate within the human auditory domain (both intensity and frequency), a rather narrow band when compared to the space available to other species, and rarely include ways to externally synchronize and trigger the delivery of auditory stimuli, with timing benchmarks highly dependent on the audio drivers used [5, 7]. High-end commercial solutions, like those provided by Tucker-Davis Technologies (TDT), can generate high-precision audio stimuli suited for neuroscience experiments [11, 9, 4] but due to their prohibitive costs have seen relatively low adoption by the experimental neuroscience community. Finally, while other open-source bespoke solutions for auditory sound generation exist [1, 5], to the best of our knowledge, choose to characterize performance at the DAC (Digital-to-Analog Converter) output instead of the amplifier and speaker element, leaving the quality metrics of the sound output partially unknown.

To meet these needs, a complete auditory stimuli delivery system was designed to provide a cost-effective open-source alternative to commercial solutions. It is comprised of a sound card device equipped with a high-precision DAC and audio amplifier, ensuring a wide-bandwidth, low-latency, audio delivery system. The flexibility of the system provides easy integration with other existing setups. At the firmware level, by implementing the Harp protocol [2] through a USB communication layer, the device can be easily controlled via Bonsai [10], or a provided graphical user interface (GUI). Adopting the Harp protocol also enables precise timestamping and synchronization with other Harp devices. Alternatively, the device can also be pre-programmed via the aforementioned GUI to be triggered through simple TTL control logic provided by an external device/microcontroller, compatible with other existing hardware ecosystems (e.g. BPOD (Sanworks), PyControl [3] or other commercially available hardware.)

Finally, the characterization of the full system (i.e. from DAC to speaker) was performed and results reporting critical specifications benchmarks are reported. Providing such characterization is critical since their acquisition might require dedicated lab equipment generally not available in neuroscience laboratories.

## 2. Hardware description

The proposed hardware system for acoustic stimulation provides an end-to-end solution. It comprises a custom-designed stereo sound card device responsible for generating high-performance analog audio signals and a low-distortion audio amplifier device for driving speakers. Both devices support wide-frequency bandwidth audio signals up to 80 kHz with a low distortion rate.

Sound waveforms can be pre-loaded in the internal memory of the sound card device, enabling low-latency sound delivery. The device also implements a high-fidelity sine-wave function generator that may be used to deliver pure tones without the need to upload a custom waveform.

An external trigger can be used to start the pre-stored sound waveform, with sub-millisecond latency, or triggered through software. In both scenarios the stimuli onset is timestamped by the microcontroller, providing the user with the real-time occurrence of this event at a resolution of 32 µs).

At the time of this manuscript, the total cost for the electronic components, printed circuit boards (PCBs) and, power supplies is 396€ (242€ sound card, 154€ audio amplifier).

The proposed system fulfills the requirements for most auditory research experiments, with the following characteristics:

- Low-cost and open-source sound system
- Sub millisecond sound stimuli latency delivery
- 192 kHz, 24 bits sound card digital-to-analog converter
- Wide frequency bandwidth system up to 80 kHz
- Low harmonic distortion
- Capable of driving speakers up to 4 W
- Compatibly with the Harp protocol, including clock synchronization between Harp devices

### 2.1 Sound card device

The sound card device is the core component of the system. It is responsible for storing and converting the digitized sound waves into high-resolution, low-distortion, and wide-band analog signals. The sound card device is composed of the power circuitry, main microcontroller and auxiliary circuitry, flash memory, audio microcontroller, audio digital-to-analog converter (DAC), and analog I/V (current-to-voltage) converter (Fig. 1).

**Figure 1:**
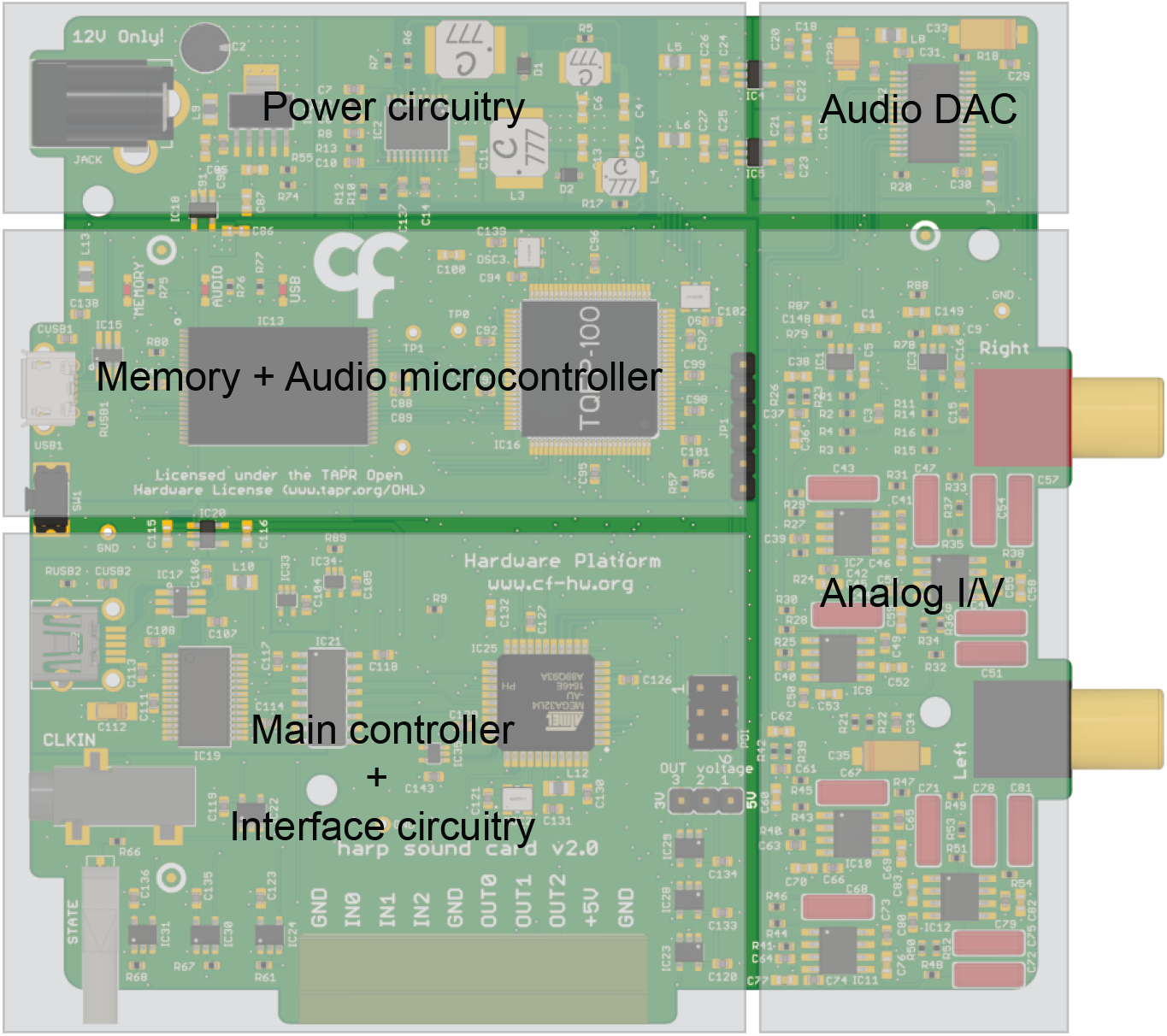
PCB rendering of the sound card depicting the main circuit blocks.

A 12 V power supply is used to power the sound card. A ±15 V step-up DC-DC followed by low-noise, low-dropout regulators are used to feed the analog I/V converter, providing a low noise and low ripple power supply to the analog circuitry. Independent analog and digital low dropout regulators are used for the audio DAC. For the digital section of the board (microcontrollers, memory and auxiliary circuitry), a 5V low-noise, fast transient-low dropout out regulator is used in series with additional 3.3 V low dropout regulators as points-of-load for different circuit parts. Given the large power dissipation of this fast-transient regulator, it is strongly recommended to attach a heat-sink to the integrated circuit.

The circuit block of the main microcontroller is responsible for the USB communication (FT232RL, FTDI), Harp clock synchronization protocol, implementation of the Harp communication protocol [2], control of the digital I/O (input/output) interface, and serial communication with the audio microcontroller. The Harp sound card is fully compatible with the Harp family of devices, enabling a seamless clock synchronization between devices guaranteeing that events from multiple Harp boards are consistently aligned.

To minimize sound latency, waveforms are stored in flash memory through a dedicated micro-USB port. The system allows sound waveforms to be loaded into the memory without disrupting potentially playing waveforms. The audio microcontroller communicates with the flash memory via a parallel bus and interfaces with the DAC audio through inter-IC sound (I2S) serial bus interface. It executes all the audio-related processing functions according to the commands received by the main controller. A reset button is available to reset the 32-bit internal processor.

The DAC audio used (AD1955, Analog Devices) was chosen because of its high-performance stereo audio digital to analog converter with a 24-bit data resolution and a sample rate of up to 192 kHz. This DAC receives an I2S input and converts it into a differential current output signal using a multi-bit Sigma-Delta modulator. The conversion of the differential current from the DAC converter is done in two stages. The initial stage, consists of an active op-amp current to voltage converter, which is used to convert each of the DAC current outputs to an analog signal. Subsequently, the I/V conversion is followed by a differential-to-single-ended buffer and low pass filter (with a cutoff frequency of 75 kHz at -3 dB), both implemented using op-amps. The sound card output voltage range is 2 V RMS.

The analog circuit block was designed to minimize noise and signal distortion, necessitating a careful choice of components. Operational amplifiers (OPA228UA, Texas Instruments) are used due to their combination of low noise and wide bandwidth operation. In the signal path, metal film chip resistors with high tolerance (*≤* 0.1 %) and low-temperature coefficient (*≤* 25 ppm), along with polypropylene film capacitors are chosen to ensure high signal generation precision and low distortion.

The sound card device has two RCA audio outputs to enable stereo audio output. The output amplitude is the typical audio line voltage, where full scale corresponds to 6 dBV (2 V RMS, 0 dBFS).

Four LEDs provide status information. The green LED will blink under the following conditions:

- Every 2 seconds during communication with a Harp-protocol host (e.g. Bonsai)
- Every 4 seconds when in standby mode
- Every 100 ms in the event of a catastrophic firmware error

The three red LEDs will provide the following information:

- Memory LED will be on when accessing the memory
- Audio LED will turn on while generating audio.
- USB LED will turn ON during USB communication or when USB communication is not available

### 2.2 Audio amplifier device

The sound card cannot provide a high-current to the output load. Hence, to drive speakers, an audio amplifier board was designed. This board can output up to 4 W of power while maintaining a low signal distortion with a flat and wide frequency bandwidth.

The primary components of the audio amplifier device consist of three key blocks: the power supply regulators and decoupling capacitors, the amplification stage, and the DC servo circuitry (Fig. 2).

**Figure 2:**
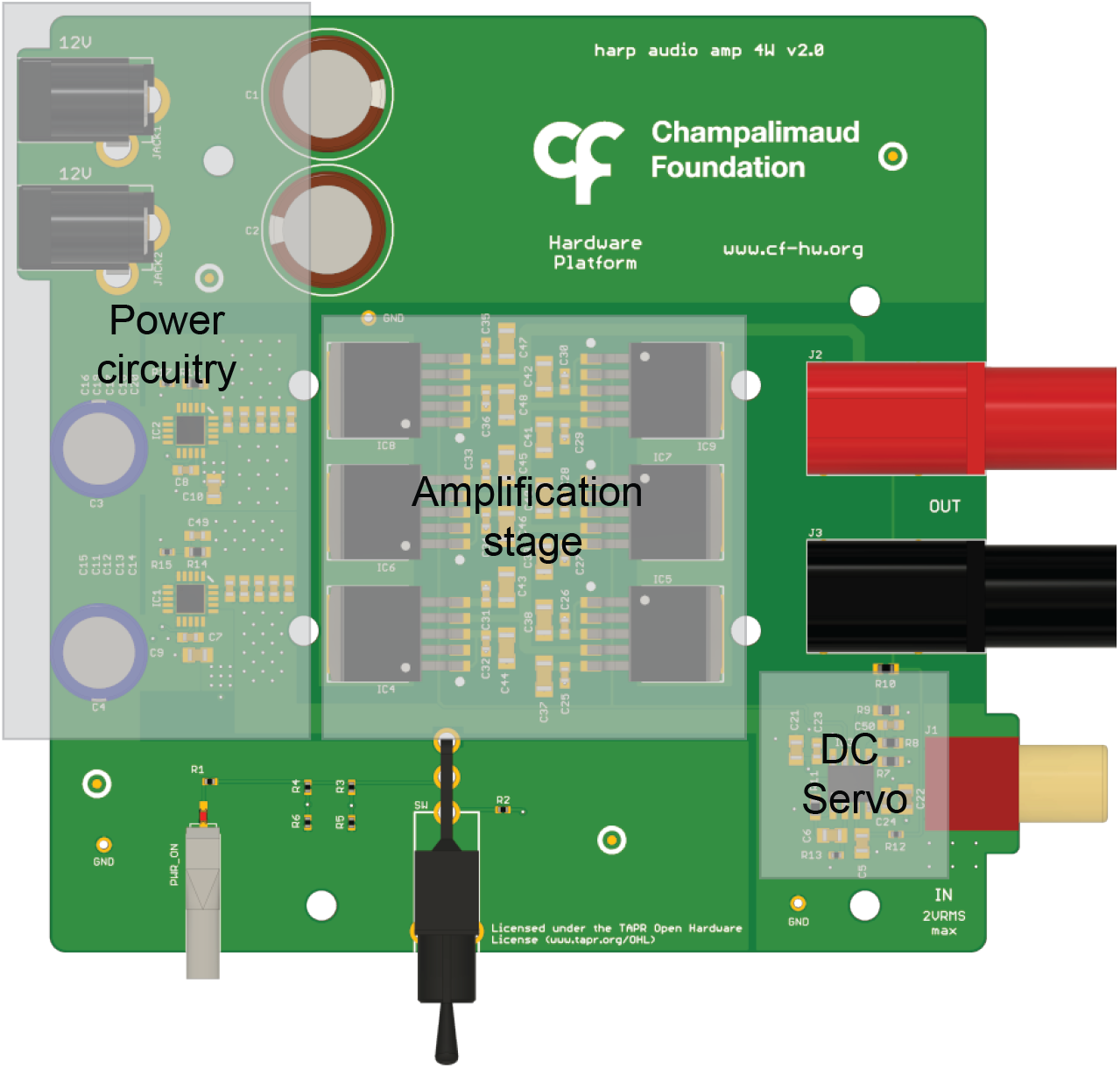
PCB rendering of the audio amplifier board depicting the main circuit blocks.

The power supply block includes two 12 V power supply connectors to generate a positive and negative voltage. This is achieved by using AC/DC power adapters with floating outputs arranged in an anti-series configuration on the board. In this configuration, the positive terminal of the top power jack is used as the +12 V supply and the negative terminal of the power jack is used as the -12 V supply voltage. The ground is connected to the remaining terminals of the power jacks. Low dropout regulators with low-output-noise and high-power supply ripple rejection are employed to supply the amplification and servo stages.

To achieve low-noise and distortion, a composite configuration was implemented. Ultra-low distortion and low noise operational amplifier (LME49720, Texas Instruments) is used, followed by a high current buffer (LME49600, Texas Instruments) operating within the feedback loop of the operational amplifier. To further increase drive current capacity, several buffers are connected in parallel. Each buffer can drive a maximum current of up to ±250 mA. Since the total noise and harmonic distortion of the buffer increase with the output power, the number of buffers employed is determined to ensure sufficient current headroom, thus keeping the distortion low.

A DC servo, consisting of an operational amplifier, compensates for the DC voltage offset in the feedback path and is implemented with a high pass filter pole set at 0.33 Hz. The role of the servo is not only to prevent a DC audio signal from being applied to the input of the amplifier device but also to remove small values of DC offset. The correction range of the offset is a function of the output voltage swing.

The amplifier is configured to operate at unity gain. Output power is specified according to the load, i.e., the impedance of the speaker that is used. For an 8 Ω load, the amplifier can deliver up to 4 W, whereas with a 4 Ω load, it can reach a maximum of 2 W (the maximum power at 4 W is limited by the current drive capacity of the amplifier, which requires a current higher than 1 A for lower loads).

In the context of the proposed audio system, given the amplifier unitary gain and the sound card maximum output voltage of 2 V RMS, the maximum output power produced is 0.5 and 1 W, at 8 and 4 Ω, respectively.

## 3. Design files

### 3.1 Design Files Summary

The design files available in the repository are listed in Tab. 1 and described below:

**Table 1:**
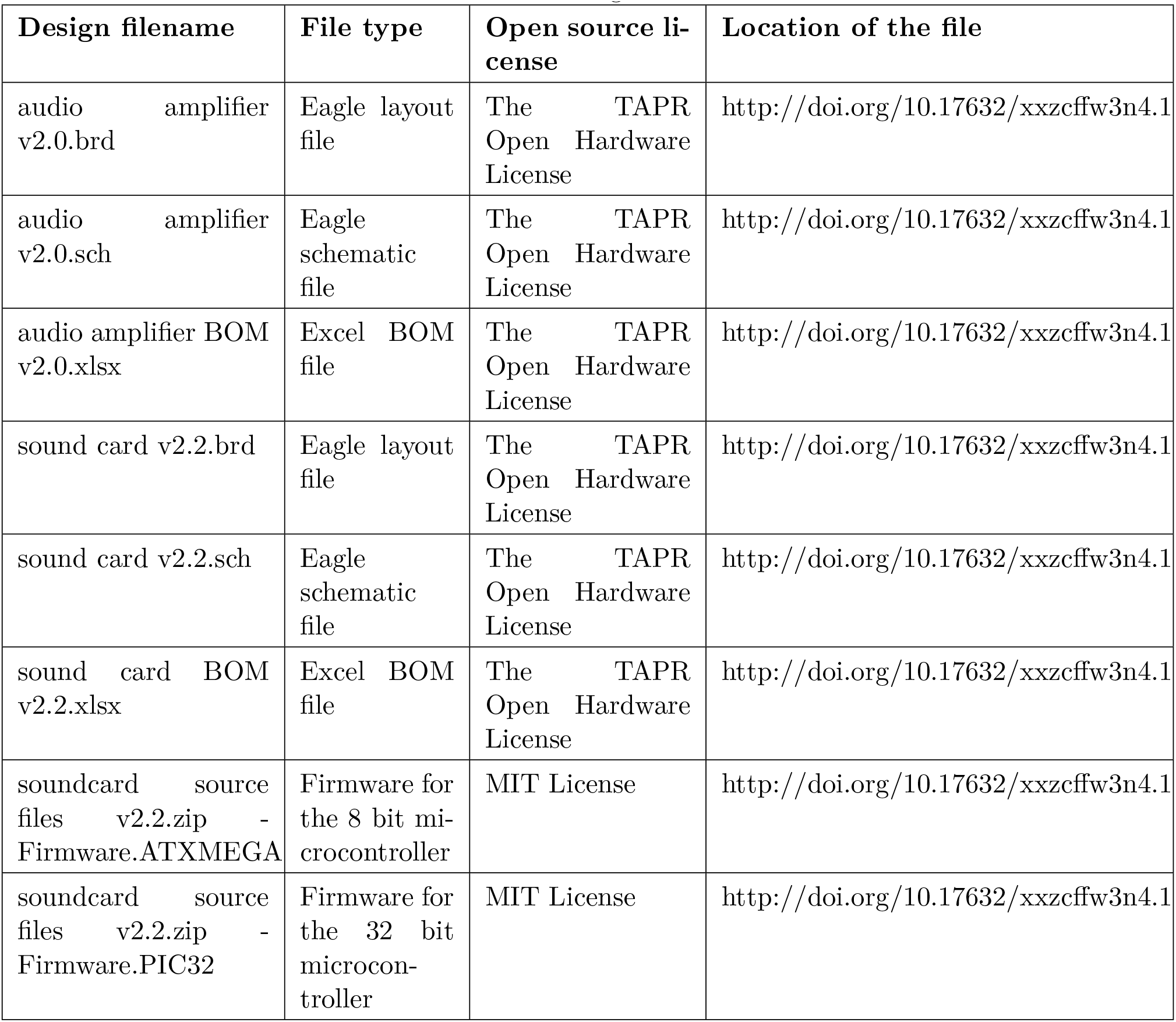
Design files list.

- audio amplifier v2.0.brd - Eagle file with the PCB layout of the audio amplifier board.
- audio amplifier v2.0.sch - Eagle file with the schematic of the audio amplifier board.
- audio amplifier BOM v2.0.xlsx - Bill of materials for the audio amplifier board, containing all the components to be assembled onto the PCB.
- sound card v2.2.brd - Eagle file with the PCB layout of the sound card.
- sound card v2.2.sch - Eagle file with the schematic of the sound card.
- sound card BOM v2.2.xlsx - Bill of materials for the sound card, containing all the - components to be assembled onto the PCB.
- Firmware.ATXMEGA - Folder with the firmware for the 8-bit processor (ATxmega128A1U, Microchip).
- Firmware.PIC32 - Folder with the firmware for the audio 32-bit processor (MZ2048EFM, Microchip).

## 4. Bill of materials

The bill of materials for both the sound and audio amplifier boards is available in the repository (http://doi.org/10.17632/xxzcffw3n4.1). At the time of this manuscript, the printed circuit boards, produced in batches of 5 at SeeedStudio, have a per-board cost of 22€ for the sound card and 19€ for the amplifier, with a total cost of 396€ for both boards. This total includes the electronic components, power supplies, cables, and PCBs, with a breakdown of 242€ for the sound card and 154€ for the audio amplifier.

## 5. Building instructions

A fully functional sound delivery system relies on the proper assembly of all the electronic components and the programming of the respective microcontrollers, as described below:

### 5.1 Printed circuit board assembly

The circuit schematic and the layout of the proposed devices were designed using the computer-aided design (CAD) software EAGLE. Users can use this software design files or the provided Gerber files to produce the PCBs. All CAD files for the device’s boards, along with the respective bill of materials and technical files required for the assembly of the boards are available in the repository (http://doi.org/10.17632/xxzcffw3n4.1).

The construction of this system requires the soldering of surface mount device (SMD) and through-hole (TH) components onto their respective PCBs (Fig. 3). Users may choose to assemble the boards themselves using hand or reflow soldering methods, provided they are comfortable with the process. Alternatively, the assembly can be outsourced to a third-party manufacturer capable of producing the PCBs, supplying the necessary components, and executing the complete assembly of the boards (e.g., PCBWay, Seeed Studio). The latter option is particularly recommended for the sound card device due to its numerous components.

**Figure 3:**
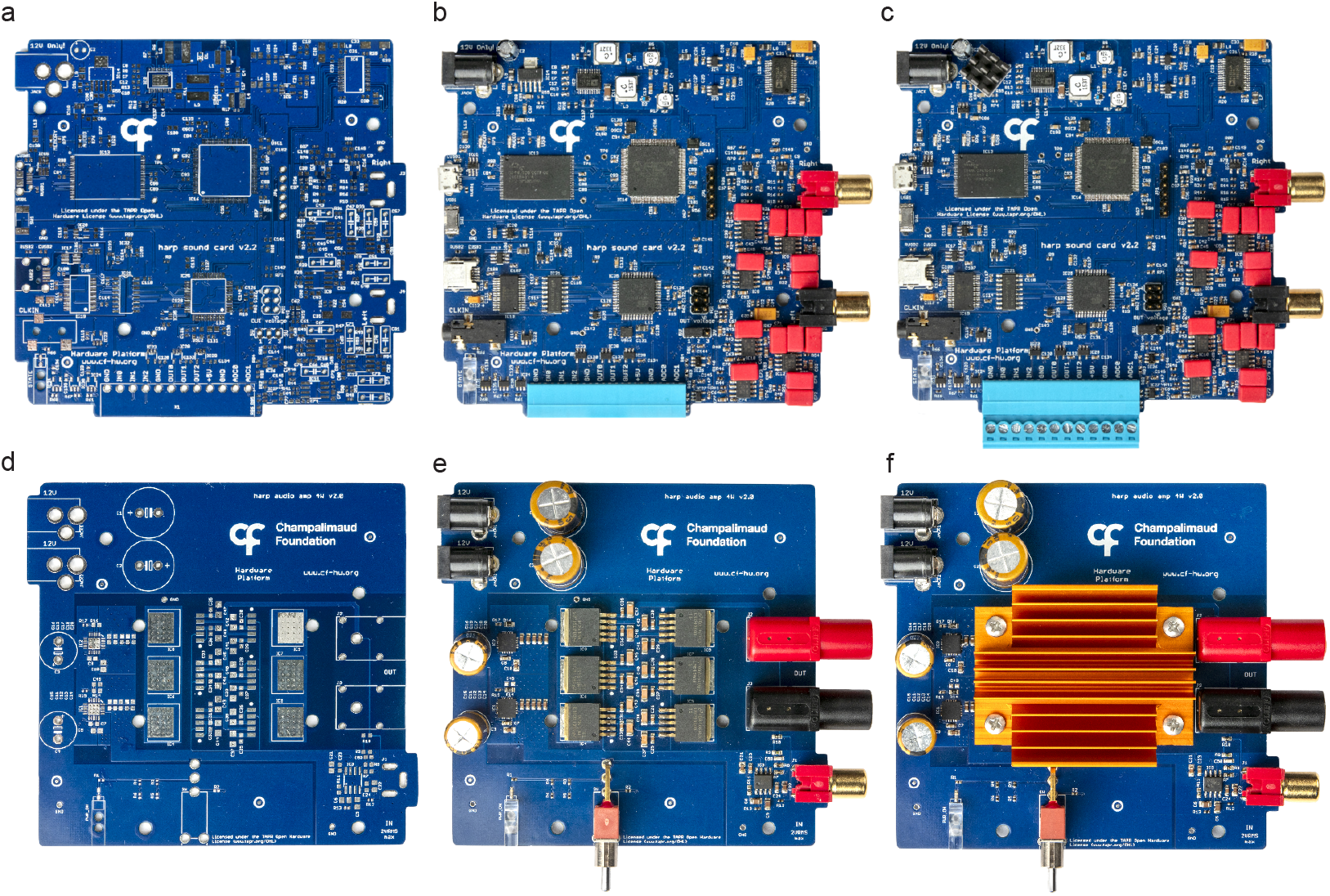
Top side view of the PCBs. a, Harp sound card PCB. b, Harp sound card assembled. c, Harp sound card fully assembled with heatsink, d, Harp audio amplifier PCB. e, Harp audio amplifier assembled. f, Harp audio amplifier fully assembled with heatsink.

### 5.2 Microcontroller programming

The sound card device requires firmware to be loaded into both microcontrollers. For each micro-controller a bootloader firmware was developed to enable subsequent firmware upgrades through the mini USB port eliminating the need for a dedicated programmer.

Therefore, it is necessary to load the respective bootloaders into the microcontrollers before uploading the firmware. The AVR XMEGA microcontroller requires the Microchip Studio IDE with and Atmel-ICE programmer or equivalent, whereas the PIC32 microcontroller uses the MPLAB IPE with a PIC32 programmer (e.g. PICKIT 3, Microchip) to load the respective bootloaders into the microcontrollers. The firmware can then be loaded using a dedicated Harp application - https://bitbucket.org/fchampalimaud/downloads/downloads/Harp_Convert_To_CSV_v1.8.3.zip.

Thus, to program the microcontroller the user needs to:

- Connect the Atmel-ICE programmer to the PDI connector in the sound card device
- Load the respective bootloader
- Connect the PIC32 programmer to the JP1 connector in the sound card device
- Load the respective bootloader
- Open the Convert to CSV application and write bootloader under List box on the Options tab
- Select the correspondent COM port and then select the firmware to be loaded for both microcontroller

## 6. Operation instructions

### 6.1 Application example

To set up the hereto-introduced audio system (Fig. 4), the initial step involves the connection of one of the sound card outputs (RCA connectors) to the amplifier input (also equipped with an RCA connector). This is the case when using a single amplification channel. A speaker can then be connected to the 4 mm banana connector outputs of the amplifier. Under no circumstance should the audio card be used to directly drive high-resistance elements like speakers.

**Figure 4:**
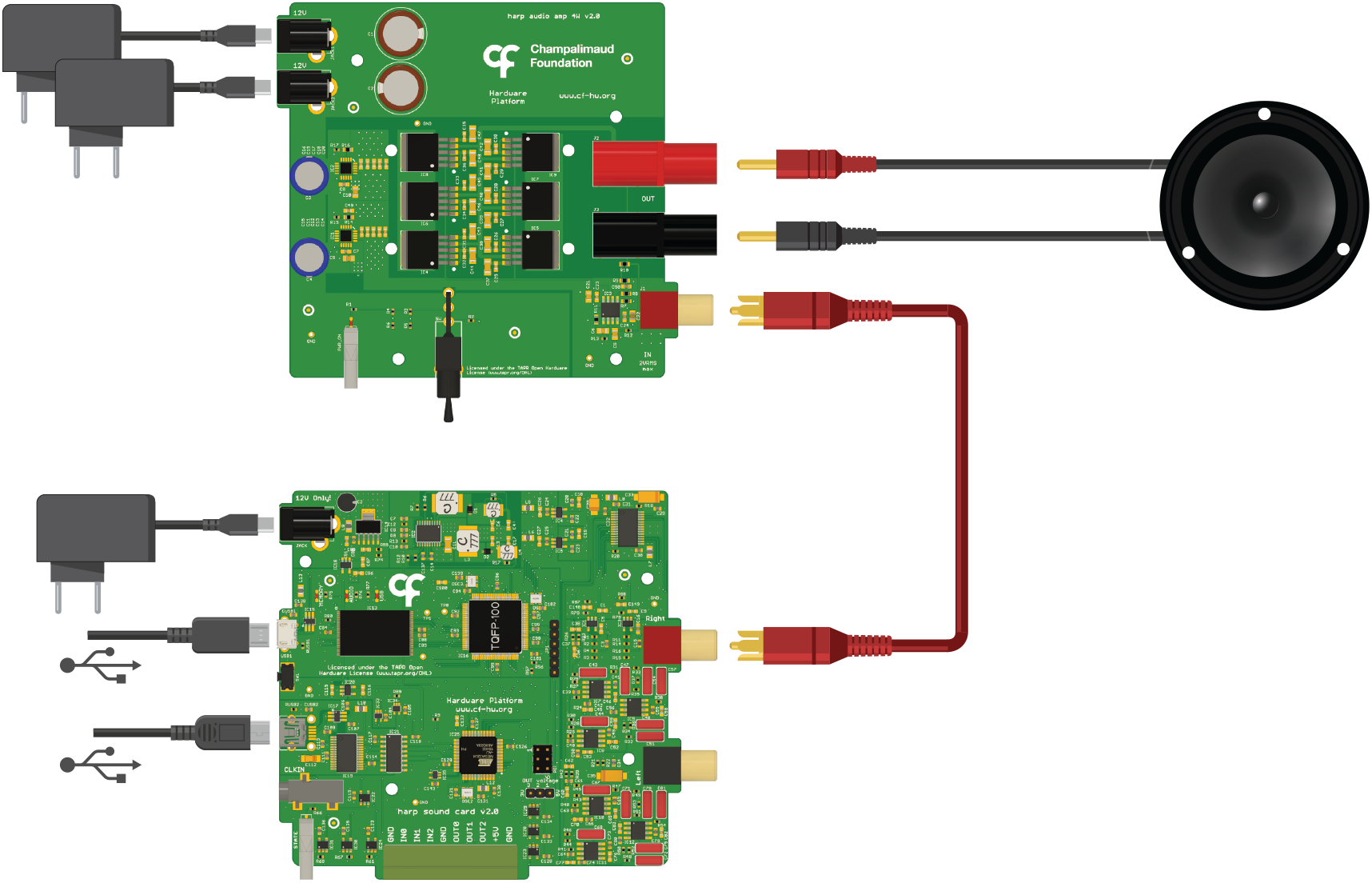
Connection diagram of proposed system – sound card device connected to the audio amplifier device where a speaker is attached.

For the power supply, three 12 V adaptors are required: two for the amplifier and one for the sound card device. It is also necessary to switch ON the power button located on the amplifier board.

A mini- and a micro-USB cable connected to a computer are needed to enable control and upload sounds to the sound card.

Finally, drivers for both USB controllers need to be installed. Instructions for installing these drivers can be found in the repository (http://doi.org/10.17632/xxzcffw3n4.1).

### 6.2 Sound card audio waveforms

The on-board sound card memory is partitioned into 32 indices (zero-indexed). The indexes from 2 to 31 are available for storing sounds, and each one of the sound indexes can store a sound file up to 8 megabytes (which corresponds to 2 million samples). An 8-MB file encodes waveforms of approximately 10.922 seconds at a 96 kHz sample rate, or 5.461 seconds at a 192 kHz sample rate (each sound sample is encoded as a 32-bit integer, although only the most significant 24 bits are used during sound reproduction). The internal memory is non-volatile and thus persists through power cycles. Example Python and Matlab scripts to generate valid waveforms are provided in the device’s repository. Users can upload waveforms via the provided GUI by connecting the micro-USB cable to the sound card device, or through Bonsai.

### 6.3 Graphical User Interface (GUI)

A graphical user interface was developed as an intuitive and easy-to-use interface to generate and upload sound files directly onto the sound card device, without the necessity for coding or the use of additional tools for sound generation (Fig. 5). This GUI was created using the graphical programming environment LabVIEW.

**Figure 5:**
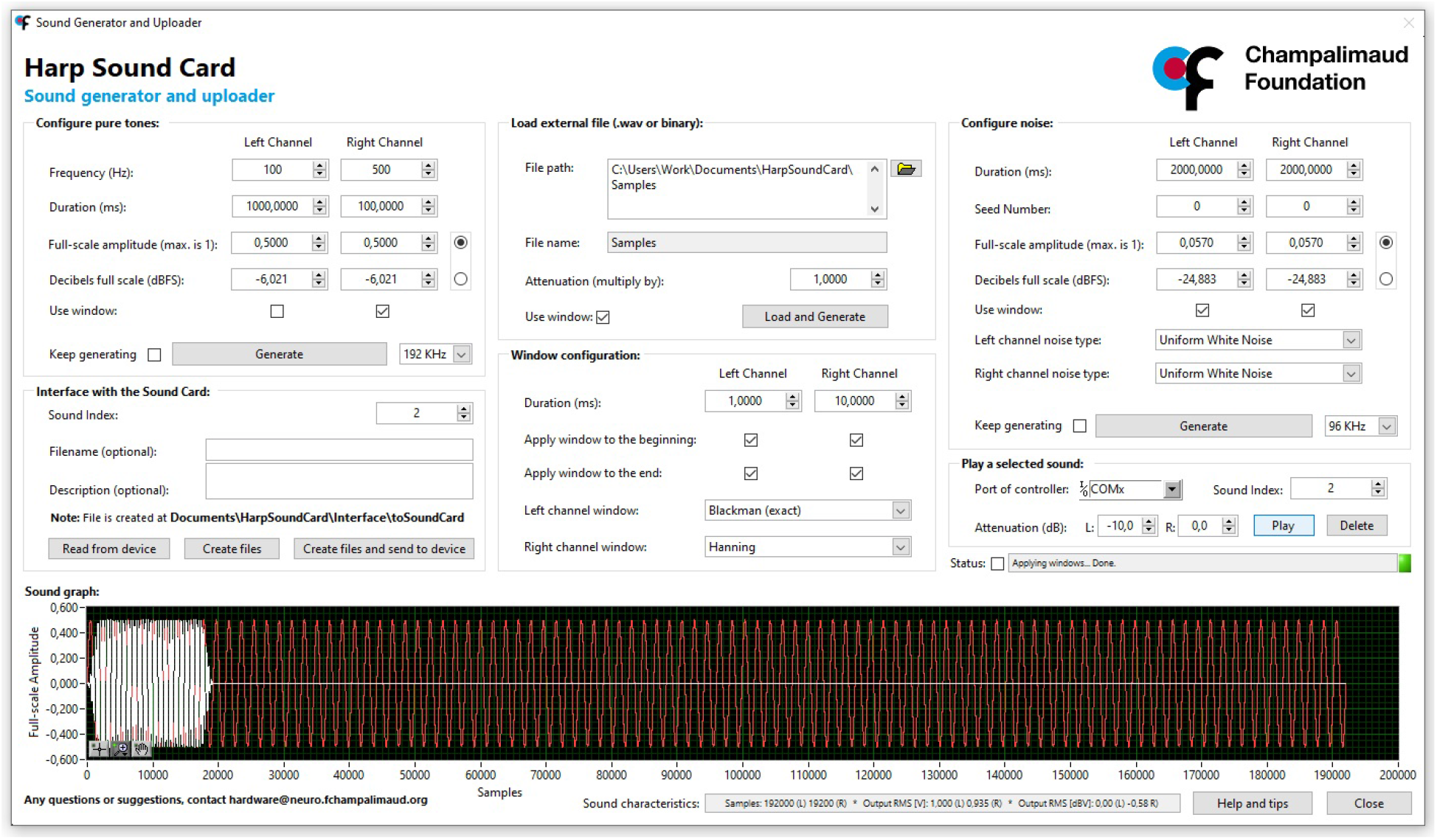
Graphical user interface screenshot.

The GUI enables the generation of pure tones with configurable frequency, duration, and amplitude for each sound card channel, as well as the generation of uniform or Gaussian white noise, with adjustable time duration and amplitude that can be loaded directly into the sound card memory. Alternatively, the GUI provides the option to open and load an external waveform (WAV or binay files) onto the board.

Whether creating a pure stone, noise or uploading an existing waveform file, users can apply a window to both of these signals (Hanning, Hamming or Blackman), with a configurable duration at the beginning, end, or both, during the generation process.

The generated signals can be previewed in the GUI in a time domain plot, depicting an amplitude versus samples, with zooming capabilities.

After the sound is generated, the user has the option to create a binary file compatible with the sound card device, which can be loaded into a specific sound index, along with an optional filename and description metadata.

### 6.4 Device interface

Various control strategies can be applied to generate sounds in the sound card device, allowing the users to select the one that is more appropriate for their experimental needs.

#### Bonsai

Bonsai is an open-source visual programming language, which provides an easy-to-use, flexible variable environment for reactive programming, tailored for data stream processing and supporting real-time interface with different hardware devices [10]. The sound card device is implemented in Bonsai through the Harp protocol [2]. Therefore, it is possible to send commands directly from the computer to the device, including playing sounds, setting attenuation values, and others. This is the most flexible and complete control option as it exposes every single function available to the board, enables precise logging of all hardware state changes, and the ability to easily interface the presented hardware with the vast ecosystem of devices already present in Bonsai (e.g. cameras, microphones, electrophysiology, etc… ).

#### GUI

In addition to the generation and uploading of waveforms, the GUI can be used to send commands to the sound card. Specifically, users can play a specific sound index with configurable attenuation, with a 0.1 dB resolution attenuation for each channel.

#### External trigger

The sound card device features a 5 V tolerant digital input that can be used to trigger a pre-selected index that contains a waveform to be played. This control strategy can be used in conjunction with either the GUI or the Bonsai to achieve reduced latency, from the trigger event to the initiation of sound.

## 7. Validation and characterization

### 7.1 Experimental validation apparatus

To fully characterize the entire signal chain and sound generation a sequential testing approach was adopted (Fig. 6). First, we characterized the performance of the sound card device (DAC). Next, we connected the audio amplifier and characterized the signal at its output with a constant external load. Finally, we recorded the sound pressure level (SPL) response of a 4 Ω speaker connected to the output of the amplifier.

**Figure 6:**
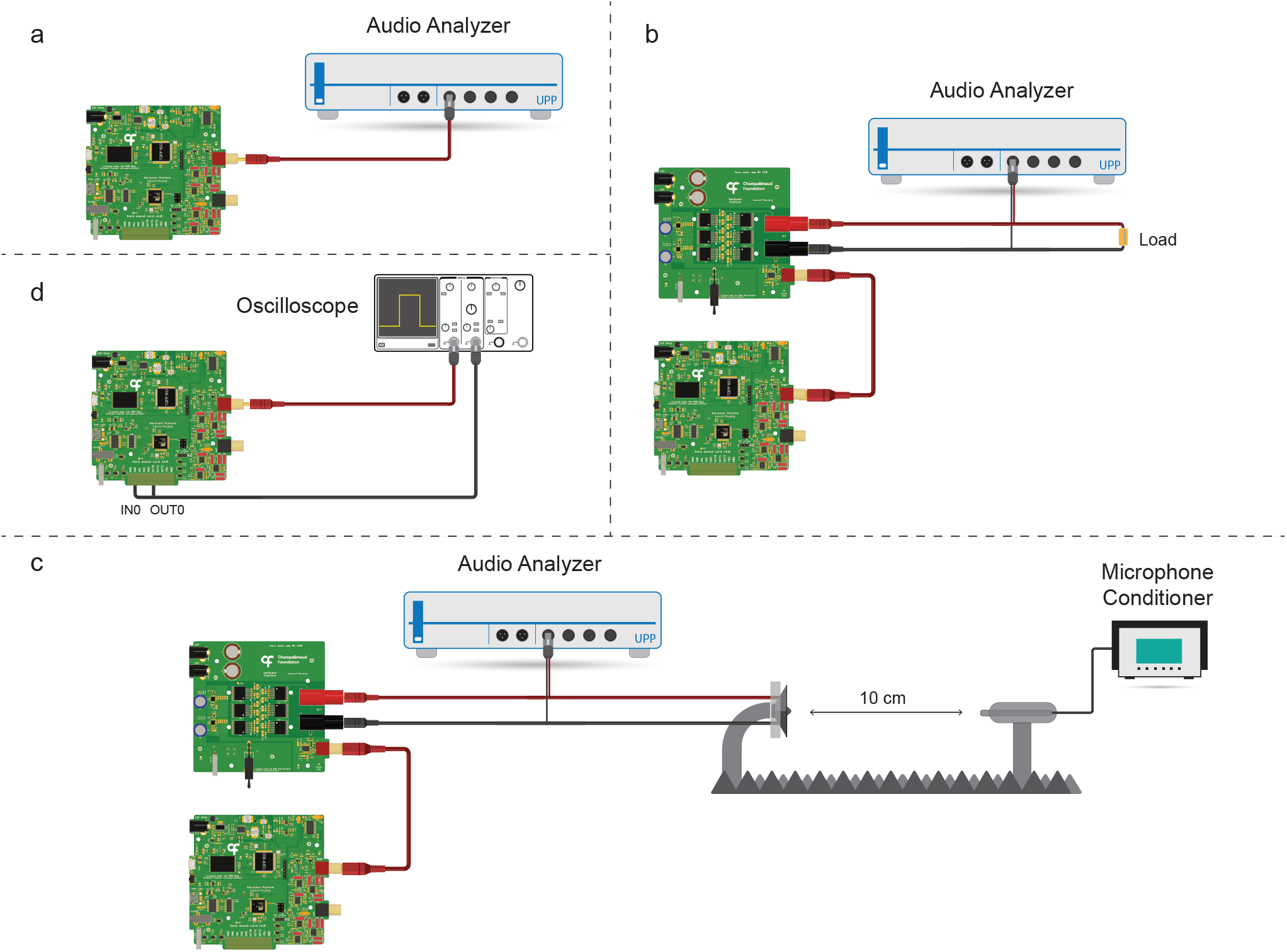
Diagram of the different experimental setups used to characterize the audio system. a, Sound card device characterization setup. b, Sound card with amplifier characterization. c, Sound system with speaker frequency response using a microphone placed 10 cm away. d, Latency measurement setup.

An industry-standard audio analyzer (Rohde & Schwarz UPP 400 Audio Analyzer) was used to measure and compute the audio metrics. To characterize the speaker response, a microphone connected to a conditioning amplifier (Brüel & Kjær conditioning amplifier and 4939-A-011 microphone) was used.

In addition to audio performance, we also measured the latencies associated with software and hardware-level triggering of waveform playback. All tests were performed by calculating the time difference between the trigger signal and an uploaded square waveform onset using an oscilloscope (Tektronix MSO 2004B).

### 7.2 Results

#### Waveform and sound output characterization

Given our sequential testing approach, we decided to report the following standard metrics [18] across all tests to facilitate performance comparison: Signal-to-noise (SNR) is defined as the ratio of the root mean square (RMS) signal amplitude to the mean value of the root-sum-square of all other spectral components, excluding harmonics and DC [8], meaning that only noise components are considered. SNR was measured by first determining the maximum amplitude of the output signal (6 dBV), followed by measuring the noise with a zero-input signal. The ratio between these two values corresponds to the SNR. The standalone sound card measures an SNR of 100 dB (113 dBA) (Fig. 7d), whereas the amplifier itself measures an SNR of 111 dB (119 dBA) (Fig. 9d).

**Figure 7:**
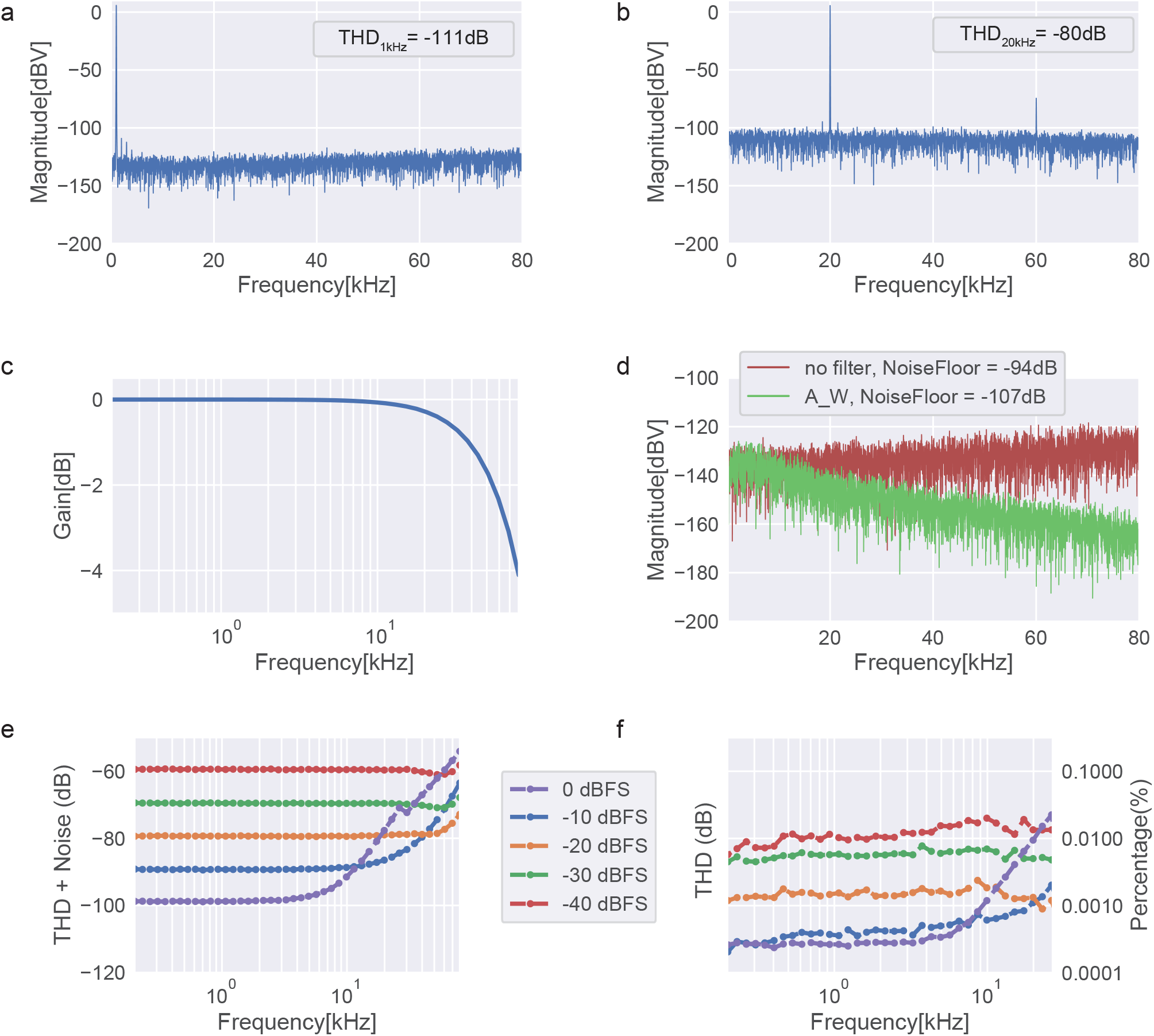
Sound card output channel, configured with a 192 kHz sampling rate. a, Fast Fourier transform (FFT) for a 1kHz sine wave signal at 0 dBFS. b, Fast Fourier transform (FFT) for a 20 kHz sine wave signal at 0 dBFS. c, Amplitude response as a function of frequency. d, Noise floor measured with zero input signal applied, with no filter and A-weighted filter for 80 kHz measuring bandwidth. e, Total harmonic distortion plus noise (THD+N) as a function of frequency for different input levels. f, Total harmonic distortion (THD) as a function of frequency for different input levels.

**Figure 8:**
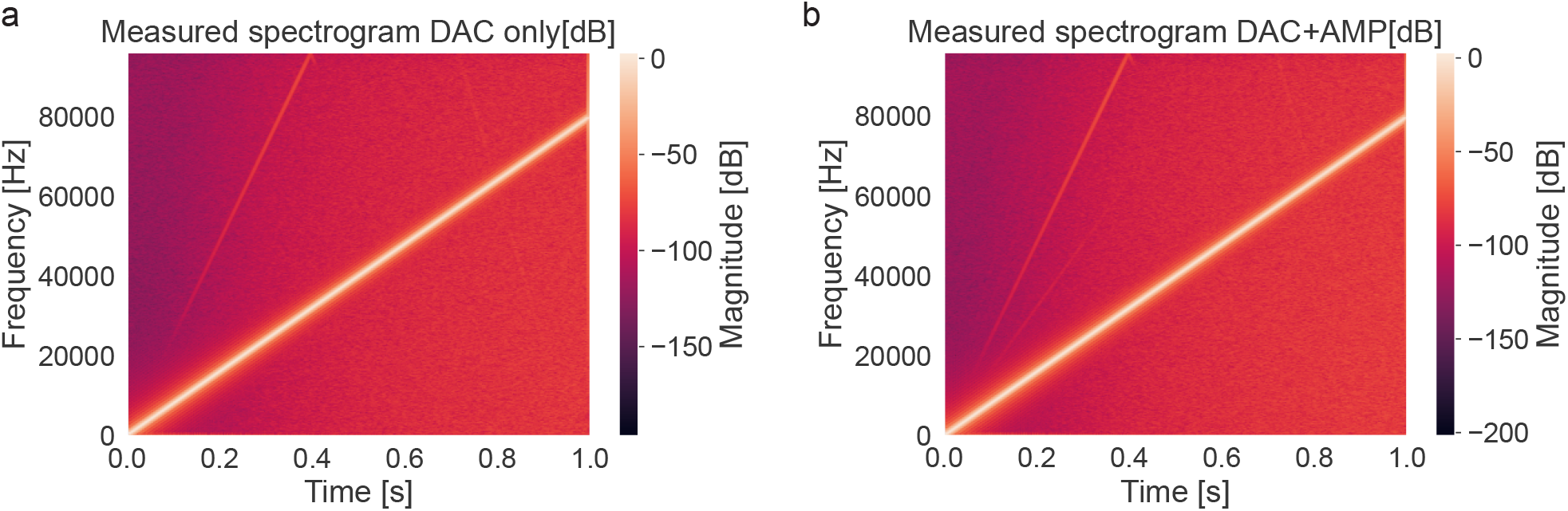
Measured spectrogram based on a 0 dBFS chirp signal with 1s duration and sweep up to 80 kHz. a, Signal measured at the output of the sound card. b, Signal measured at the output of the audio amplifier connected to the sound card.

**Figure 9:**
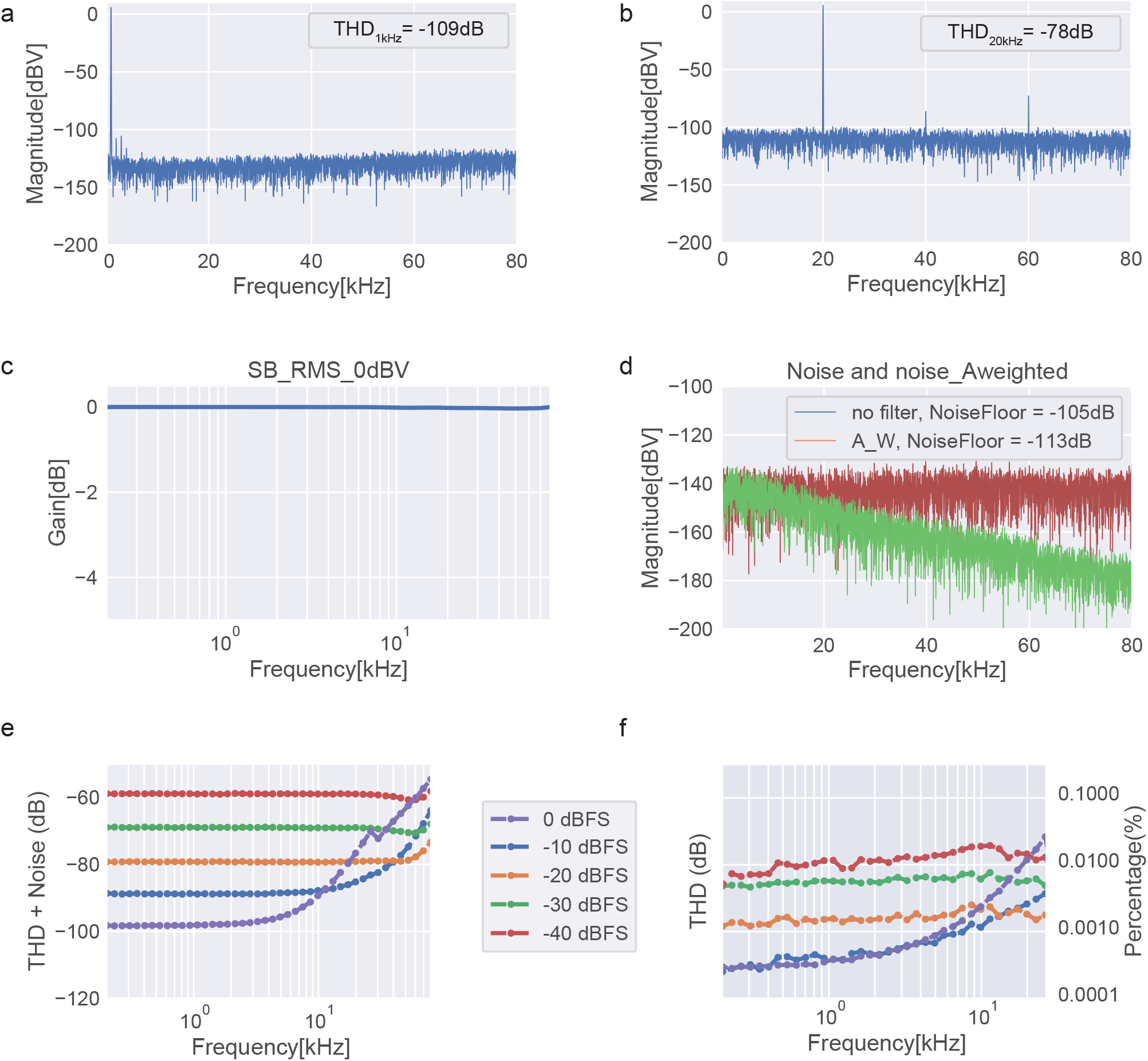
Output channel of the sound card and amplifier path with a 4 Ω load, configured with a 192 kHz sampling rate. a, Fast Fourier transform (FFT) for a 1kHz sine wave signal at 0 dBFS. b, Fast Fourier transform (FFT) for a 20 kHz sine wave signal at 0 dBFS. c, Amplifier amplitude response as a function of frequency. d, Noise floor measured with zero input signal applied, with no filter and A-weighted filter for 80 kHz measuring bandwidth (amplifier only). e, Total harmonic distortion plus noise (THD+N) as a function of frequency for different input levels. f, Total harmonic distortion (THD) as a function of frequency for different input levels.

The amplitude-frequency responses for both the sound card and the amplifier were measured individually (Fig. 7c and 9c). The sound card has a low pass filter with a pole at 75 kHz whereas the amplifier shows a flat response across the entire bandwidth (*<* 0.1 dB variation over the full bandwidth).

To assess the audio distortion of the system, total harmonic distortion (THD) and total harmonic distortion plus noise (THD+N) metrics are reported. THD is defined as the ratio of the root mean square value of the fundamental signal to the mean value of the root-sum-square of its correspondent harmonics measured in the specified bandwidth [18]. In other words, THD quantifies the “purity” of the sound generated. THD+N considers also the noise components of the signal. Fig. 7e,f and Fig. 9e,f depict the frequency response of THD and THD+N, respectively, for both the standalone sound card and the sound card paired with the amplifier, using an external load (4 Ω) across different input signal levels. A low resistance of 4 Ω was chosen to induce a higher current draw, representing a worst-case scenario in terms of distortion rates in the output signals.

To summarize the response of the system across the full frequency bandwidth, we constructed a chirp signal with 1 s duration linearly sweeping up to 80 kHz, and played it through the sound card. The resulting spectrograms were captured at the output of the sound card, and then at the output of the audio amplifier when connected to the sound card under a constant load. Harmonic distortion values remained qualitatively similar between the two conditions (Fig. 8). Nonetheless, a residual distortion component relative to the second harmonic of the signal can be detected, in addition to pre-existing the 3rd harmonic distortion of the signal.

As a final test, a 4 Ω speaker (Tymphany XT25SC90-04) was connected to the output of the amplifier board, and a frequency response analysis was performed by generating and playing 100 distinct pure tones sweeping from 200 Hz to 80 kHz at an amplitude of -20 dBFS. A microphone connected to a conditioning amplifier (Brüel & Kjær conditioning amplifier and 4939-A-011 microphone) aligned with the speaker positioned at a 10 cm distance was used to record the SPL.

The results are depicted in Fig. 10, showing a variation response of 35 dB over the 80 kHz measuring bandwidth. It is important to note that these variations likely result from speaker-specific distortion. Nevertheless, we also show that, should a specific experiment require a flat response curve across all frequencies, a simple calibration procedure can be performed by selectively modulating the intensity of the different pure tone frequencies. Specifically, considering the magnitude of the SPL of the non-calibrated sweep, the amplitude of the 100 pure tones was adjusted to get a 70 SPL flat response. The calibrated response exhibits a variation of ±1.1 dB over the 200 Hz - 80 kHz range.

**Figure 10:**
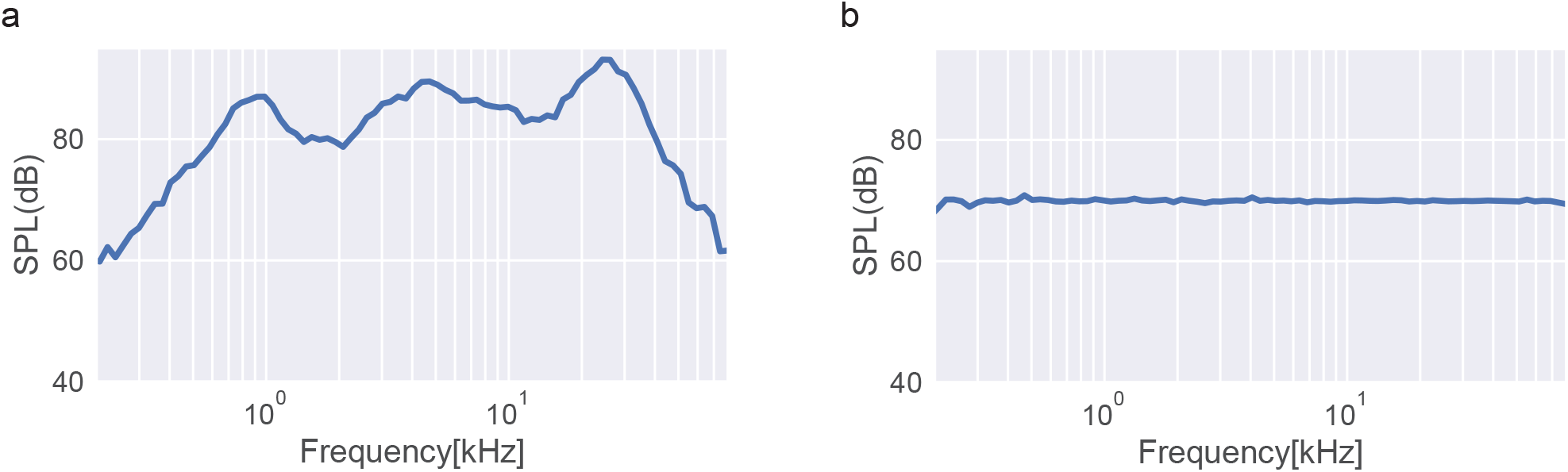
Frequency response at a distance of 10 cm from a 4 Ω speaker with a -20 dBFS input signal. a, Frequency response as a function of Sound Pressure Level (SPL). b, Frequency response as a function of SPL after calibration to 70 dB.

#### Characterizing trigger latency

Given the tight timing often required for behavior and neurophysiological experiments, the current system affords both hardware and software triggers to instantiate a waveform playback.

As mentioned above, we measured the latency between an external TTL trigger and the onset of a previously uploaded square wave using the configuration illustrated in Fig. 6d. The protocol was repeated 100 times and the results are summarized in Fig. 11b. Overall, all triggers resulted in a sub-millisecond latency (698 ± 104 µs, mean ± std), an adequate benchmark for most neuroscience behavior and neurophysiological experiments.

**Figure 11:**
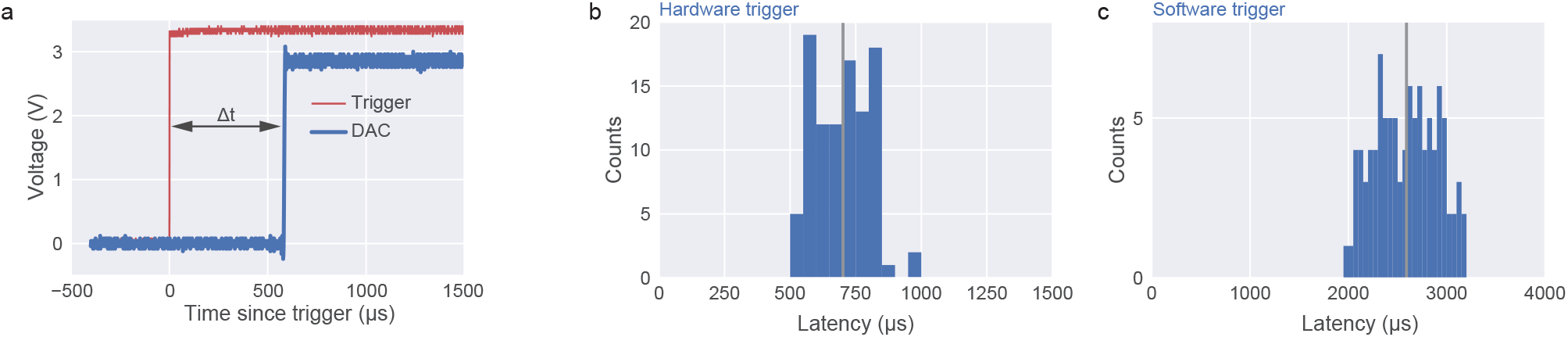
Triggered playback latency. a, Trigger (blue) and generated square pulse signal (red) traces. b, Histogram for latencies measured from 100 trials with an external hardware trigger. c, Same as b, for a software trigger. See text for further details.

As an alternative, sound playback can be triggered via software. Such an option will sacrifice latency but afford users greater flexibility. Since one-way delays are notoriously difficult to calculate (as the host runs on a different clock), we opted to instead calibrate what is usually known as a round-trip delay. Briefly, a command was sent to trigger a digital output shorted to a digital input in the same board. Once the computer host received the event corresponding to this digital input toggle, a message was immediately issued to start the audio playback. The reported latency will thus correspond to the sum between the two events: “Sound card digital input event → PC” plus “PC → playback command”. Assuming symmetric latencies the one-way delay should be roughly the measured amount (2584 ± 313 µs) (Fig. 11c).

Tab. 2 and Tab. 3 detail the most relevant technical specifications for the sound card and audio amplifier device, respectively.

**Table 2:** Sound card device specifications.

| Specification | Value |
| --- | --- |
| SNR | 100 dB — 113 dBA (20 Hz – 80 kHz @ 2 V rms) |
| THD | -109dB (1 kHz @ 2 V rms) |
| Noise Floor | 20 μV rms — -94 dB (20 Hz – 80 kHz) |
| Maximum sampling rate | 192 kHz |
| Number of channels | 2 channels) |
| Number of bits | 24 bits |
| Output Voltage | 2 V rms |

**Table 3:**
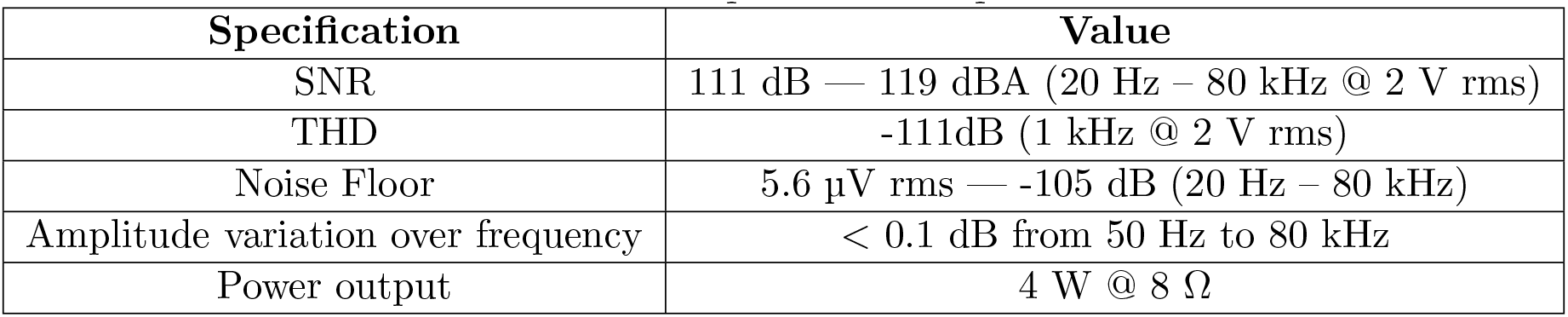
Audio amplifier device specifications.

### 7.3 Conclusions

In summary, the proposed audio system not only offers an open-source cost-effective alternative to commercial solutions for high-performance acoustic stimulation, especially in experiments requiring low audio distortion, but also presents a solution with reduced onset sound latency. Furthermore, this system has an extended bandwidth range of up to 80 kHz, surpassing most of the audio systems available that are designed for the human hearing frequency bandwidth range. Both the sound card and audio amplifier devices can be paired in the same audio system or used independently with other equipment.

In addition to the hardware development, a thorough characterization of the audio devices and the entire system up to the sound delivery was conducted. This is particularly important as most technical documentation for audio products relies on a limited set of flagship metrics, valid only under specific conditions, often lacking the performance analysis for a broader range of usage scenarios, namely the frequency response over the entire bandwidth or with different input signal ranges.

## 8. CRediT author statement

**Artur Silva**: Conceptualization, Methodology, Validation, Writing - Original Draft, Writing - Review & Editing, Visualization. **Filipe Carvalho**: Conceptualization, Methodology, Software, Validation, Writing - Review & Editing. **Bruno Cruz**: Formal analysis, Software, Writing - Review & Editing, Visualization.

## 9. Acknowledgements

We thank Dario Bento for assembling the boards, Mafalda Valente for providing support with the microphone related measurements, Gon çalo Lopes for help with the Bonsai interface, Alexandre Azinheira for the boards photos and Hugo Marques for proofreading the manuscript.

This research did not receive any specific grant from funding agencies in the public, commercial, or not-for-profit sectors.

